# Molecular imaging with aquaporin-based reporter genes: quantitative considerations from Monte Carlo diffusion simulations

**DOI:** 10.1101/2023.06.09.544324

**Authors:** Rochishnu Chowdhury, Jinyang Wan, Remy Gardier, Jonathan Rafael-Patino, Jean-Philippe Thiran, Frederic Gibou, Arnab Mukherjee

## Abstract

Aquaporins provide a new class of genetic tools for imaging molecular activity in deep tissues by increasing the rate of cellular water diffusion, which generates magnetic resonance contrast. However, distinguishing aquaporin contrast from the tissue background is challenging because water diffusion is also influenced by structural factors such as cell size and packing density. Here, we developed and experimentally validated a Monte Carlo model to analyze how cell radius and intracellular volume fraction quantitatively affect aquaporin signals. We demonstrated that a differential imaging approach based on time-dependent changes in diffusivity can improve specificity by unambiguously isolating aquaporin-driven contrast from the tissue background. Finally, we used Monte Carlo simulations to analyze the connection between diffusivity and the percentage of cells engineered to express aquaporin, and established a simple mapping that accurately determined the volume fraction of aquaporin-expressing cells in mixed populations. This study creates a framework for broad applications of aquaporins, particularly in biomedicine and in vivo synthetic biology, where quantitative methods to measure the location and performance of genetic devices in whole vertebrates are necessary.

## INTRODUCTION

Genetically encoded reporters are essential tools for monitoring molecular signals in living systems. In synthetic biology, reporters based on fluorescent and bioluminescent proteins provide a natural approach for measuring and optimizing the performance of genetic systems^1^. However, optical reporters are of limited use for tracking genetically engineered devices in living animals due to absorption and scattering of light in thick tissue^2–4^. Unlike optical methods, magnetic resonance imaging (MRI) can image deep tissues and generate volumetric scans with a high spatial resolution. We recently developed an MRI-based reporter that enables imaging of genetic activity in deep tissues^5–7^. This reporter utilizes aquaporin-1 (Aqp1), a channel that allows water molecules to diffuse freely across the plasma membrane^8,9^. In contrast to wild-type cells, which restrict water movement owing to the low permeability of the plasma membrane, cells engineered to express Aqp1 allow the free exchange of water (**Fig. 1a**). Accordingly, Aqp1 expression increases the molecular diffusivity of water in cells and tissues, which can be visualized using an MRI technique known as diffusion-weighted imaging^10,11^. In this technique, pulsed magnetic field gradients create a phase dispersion in water molecules, producing a signal that decays in proportion to water diffusivity.

**Figure 1:**
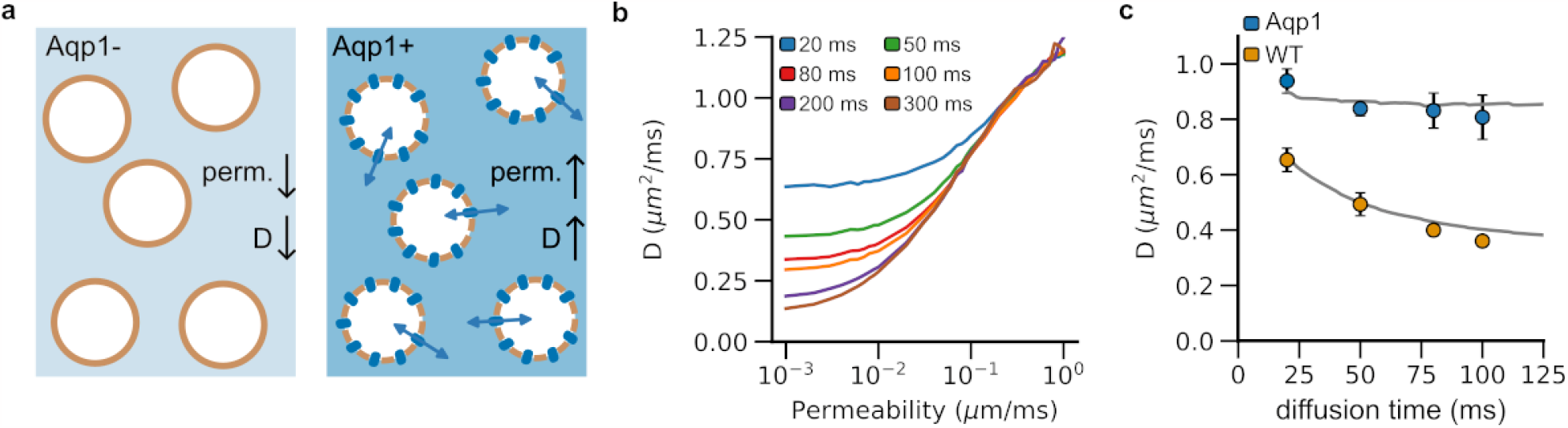
Monte Carlo simulations of water diffusion in cells with varying plasma membrane permeability. **a**, Engineering cells to express Aqp1 makes their membrane more permeable to water molecules than wild-type (viz. non-engineered) cells. **b**, Molecular diffusivity (*D*) increases with membrane permeability. Longer diffusion times lead to a decrease in diffusivity as a larger number of spins encounter the plasma membrane, which restricts free movement of water molecules. **c**, Diffusivity decreases more markedly with diffusion time in wild-type cells (less permeable) than in cells engineered to express Aqp1 (more permeable). Therefore, extended diffusion times (∼ 100 *ms*) are required to maximize Aqp1-driven contrast. The solid lines represent the simulated diffusivities for a synthetic substrate consisting of spherical cells of radius 7.6 μ*m* packed to yield a total intracellular volume fraction of 0.65. Wild-type and Aqp1-expressing cells were modeled using permeability coefficients of 0.012 and 0.138 μ*m*/*ms*, respectively. Circles denote experimental data obtained from pellets of CHO cells at 7 T. Error bars represent the standard deviation (*n* ≥ 5 biological replicates).

Although Aqp1 provides a promising tool for monitoring genetic systems using MRI, variations in tissue microstructure, such as cell size and intracellular volume fraction, can affect tissue water diffusion^12–17^, thereby making it difficult to unambiguously link Aqp1-driven signals with a specific genetic output or cell-type. For example, a decrease in intracellular volume fraction as a result of cell death or apoptotic shrinkage could lead to an increase in the rate of water diffusion in tissues independent of Aqp1 expression^18–20^. Conversely, an increase in cell size owing to swelling or mitotic growth arrest can decrease the rate of water diffusion^12,21,22^. To expand Aqp1 into a broadly useful reporter for deep-tissue imaging, we need a mechanistic framework that predicts how changes in molecular diffusivity induced by Aqp1 expression are affected by cell radius, packing density, and the volume fraction of Aqp1-expressing cells.

Monte Carlo diffusion simulations, which compute the Brownian motion of water molecules in the presence of a dephasing magnetic field gradient, are widely used to investigate the correlation between molecular diffusion of water and tissue morphology^23–37^. For example, Monte Carlo simulations have been used to quantify changes in white matter diffusivity caused by the swelling and beading of neurites during ischemic stroke^33,38^. In cancer biology, Monte Carlo methods have been used to explore the effects of cell size, packing density, and compartment volume fractions on tumor diffusion^27,32^. We recently applied Monte Carlo diffusion simulations to generate ground-truth diffusion datasets, which we used to compare the accuracies of various analytical models for estimating tissue microstructure using diffusion-weighted MRI^36^.

In this study, we developed and experimentally validated a Monte Carlo simulator to model water diffusion in cells engineered to express Aqp1. We showed that Aqp1 operates as an effective reporter over a wide range of cell sizes and volume fractions, driving larger changes in molecular diffusivity than those seen in wild-type (viz. non-engineered) cells. We also identified the range of cellular radii and volume fractions that lead to nonspecific enhancements in molecular diffusivity, thereby making it nontrivial to unambiguously discern Aqp1-based MRI signals from the tissue background. We further show that the time-dependence of diffusion coefficients can be exploited to specifically image Aqp1-expression without interference from the background. Finally, we used Monte Carlo simulations to analyze the correlation between diffusivity and the volume fraction of Aqp1-expressing cells and demonstrated that a simple log-linear model was sufficient to measure Aqp1-expressing cells in mixed-cell populations, thereby combining cell-type specificity with quantitative imaging.

## RESULTS

### Aqp1 generates a substantial increase in diffusivity at long diffusion times

We designed our computational tissue phantom to consist of spherical cells (7.6 μ*m* radius) tightly packed to yield an intracellular volume fraction of 0.65. This configuration mimics our experimental system comprising lightly centrifuged pellets of Chinese hamster ovary (CHO) cells. We used Monte Carlo simulations to compute diffusivities for a range of permeability coefficients and diffusion times. Consistent with previous studies of water diffusion in similar geometries^39^, the diffusivity increases with membrane permeability, rising sharply as the permeability crosses ∼10^−2^ μ*m*/*ms* (**Fig. 1b**). Longer diffusion times lead to a decrease in diffusivity as more spins encounter the plasma membrane, which restricts water movement (**Fig. 1b,c**). In contrast, permeable membranes (e.g., due to Aqp1 expression) do not substantially hinder the free movement of water molecules and thus the time dependence of diffusion become less pronounced with increasing permeability (**Fig. 1c**). Accordingly, extended diffusion times are optimal for maximizing diffusion-weighted contrast induced by Aqp1 expression.

Next, we used diffusion-weighted MRI to measure diffusivities in pellets of both wild-type cells and cells in which the Aqp1 reporter was introduced as a transgene using lentiviral transduction (**Supplementary Fig. 1**). Our experimental estimates agreed with diffusion coefficients computed from Monte Carlo simulations where we modeled wild-type and Aqp1-CHO cells using permeability coefficients of 0.012 and 0.138 μ*m*/*ms* respectively based on previously published estimates^40,41^ (**Fig. 1c**). Notably, at a diffusion time of 100 *ms*, Aqp1-expressing cells showed a 124 ± 14 % (mean ± s.d., *n =* 6) increase in diffusivity compared to wild-type cells, which aligns well with the 112 % increase predicted from our simulations. Longer diffusion times will further enhance the Aqp1-driven increase in diffusivity, though this also decreases the signal-to-noise ratio in diffusion-weighted images. Accordingly, in the remainder of the study, we chose 100 *ms* as an optimal diffusion time to characterize the performance of Aqp1 in various tissue configurations.

### Changes in tissue microstructure may elevate diffusion rates, at times overlapping with Aqp1-driven changes in molecular diffusivity

We applied the Monte Carlo diffusion framework to explore the effects of tissue microstructure parameters, namely cell size (*r*) and intracellular volume fraction (*v*_*f*_) on the diffusivities of wild-type and Aqp1-expressing CHO cells. Our specific goal was to identify cell sizes and volume fractions that would elevate the rate of water diffusion and by doing so create the same effect on MRI readouts as the Aqp1 reporter (**Fig. 2a**). In practical terms, this parameter space represents tissue configurations where Aqp1 signals are hard to tell apart from the tissue background. For a fixed extracellular volume fraction (*v*_*f*_ = 0.65), we found that wild-type diffusivity was highly sensitive to cell size, increasing by as much as 122 % as the radius was varied from 5 μ*m* to 25 μ*m* (**Fig. 2b**). Expression of Aqp1 reduced the cell size dependence, producing a 36 % increase in diffusivity over the same size range (**Fig. 2b**). Wild-type cells were also more sensitive to changes in the intracellular volume fraction and their diffusivity increased by 177 % (compared to 80 % for Aqp1 cells) when the volume fraction was increased over a 5-fold range, while keeping the cell radius fixed (**Fig. 2c**).

**Figure 2:**
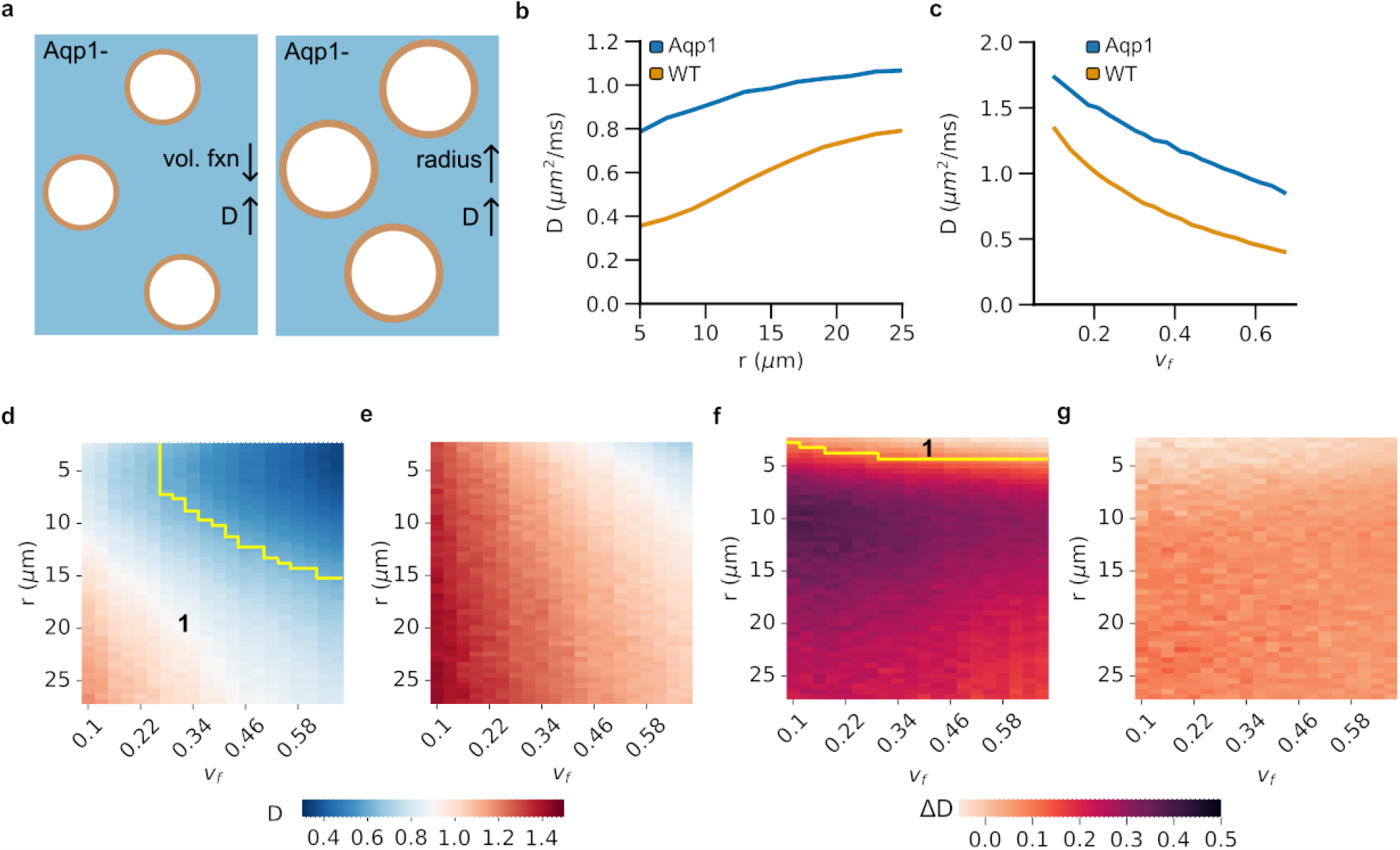
Effect of cell size and intracellular volume fraction on diffusivity. **a**, Changes in tissue microstructure, such as reduced intracellular volume fraction (*v*_*f*_) or larger cell radius (*r*), can increase diffusivity (*D*) similar to Aqp1 expression. **b**, Simulated diffusivity at a fixed volume fraction (*v*_*f*_ = 0.65) increases with cell radius in both Aqp1-expressing and wild-type cells, but Aqp1 expression reduces the size dependence of diffusivity. **c**, Simulated diffusivity for a given cell size (*r* = 7.6 μ*m*) shows an inverse correlation with intracellular volume fraction in Aqp1-expressing and wild-type cells. **d**, Heatmap showing the combined dependence of diffusivity on cell size and intracellular volume fraction in wild-type and **e**, Aqp1-expressing cells. In the region marked as 1, Aqp1 signals are difficult to distinguish from the tissue background because wild-type diffusivities fall within the same range of diffusivities observed in Aqp1-expressing cells. **f**, Heatmap showing the dependence of differential diffusivity, viz. Δ*D* = *D*_20 *ms*_ − *D*_100 *ms*_, on cell size and intracellular volume fraction in wild-type and **g**, Aqp1-expressing cells. The Δ*D* metric significantly reduces the overlap (region of overlap marked as **1** in the heatmap) between cell populations expressing Aqp1 and wild-type cells, thereby providing a reliable readout of Aqp1 expression that is unaffected by tissue microstructure parameters. The color-bars represent molecular diffusivity of water (*D*) or differential diffusivity (Δ*D*) in units of *um*^2^/*ms*. Wild-type and Aqp1-expressing cells were modeled with permeability coefficients of 0.012 and 0.138 μ*m*/*ms*, respectively.

To map the parameter space where Aqp1 signals are potentially masked by elevations in background tissue diffusion, we calculated diffusivities while simultaneously varying both the intracellular volume fraction and cell size (**Fig. 2d,e**). In general, configurations with large cells and low intracellular volume fractions had diffusivities high enough to overlap with Aqp1 signals (**Fig. 2d,e**). We wondered whether we could use the distinct time-dependence of diffusivity in wild-type and Aqp1-expressing cells (**Fig. 1c**) to distinguish between signals resulting from Aqp1 expression and nonspecific diffusion enhancements caused by changes in tissue microstructure. Notably, in sharp contrast to Aqp1-expressing cells, the diffusivity of wild-type cells changes rapidly with diffusion time (**Fig. 1c**). Accordingly, we explored how the difference in simulated diffusivities between 20 *ms* and 100 *ms* (Δ*D* = *D*_20_ − *D*_100_) changes as a function of radii and intracellular volume fractions (**Fig. 2f,g, Supplementary Fig. 2**). Strikingly, the Δ*D* metric successfully differentiated between wild-type and Aqp1-expressing cells for nearly all combinations of radii and volume fractions tested (**Fig. 2f**), indicating that difference imaging at two diffusion times provides a unique approach for accurately identifying expression of the Aqp1 reporter regardless of changes in microstructure parameters.

### Monte Carlo simulations allow mapping of Aqp1 volume percentage in mixed cell populations

In many applications of reporter gene technology, such as tracking cell therapies and monitoring transcriptional activity, only a subset of cells may express the reporter at a given time. In these scenarios, the ability to quantify the fraction of reporter-expressing cells permits a richer description of the underlying biological process. To this end, we hypothesized that the dependence of molecular diffusivity on the volume percentage of Aqp1-cells (*v*_*Aqp*1_) in a mixed population could be used for quantitative imaging of reporter gene expression (**Fig. 3a**). To test this idea, we analyzed the relationship between diffusivity calculated by our Monte Carlo simulations and *v*_*Aqp*1_ in mixed populations comprising CHO-Aqp1 cells interspersed with wild-type cells in varying ratios. We found that a simple log-linear function quantitatively describes the dependence of molecular diffusivity on *v*_*Aqp*1_ (**Fig. 3b, Supplementary Fig. 4**).

**Figure 3:**
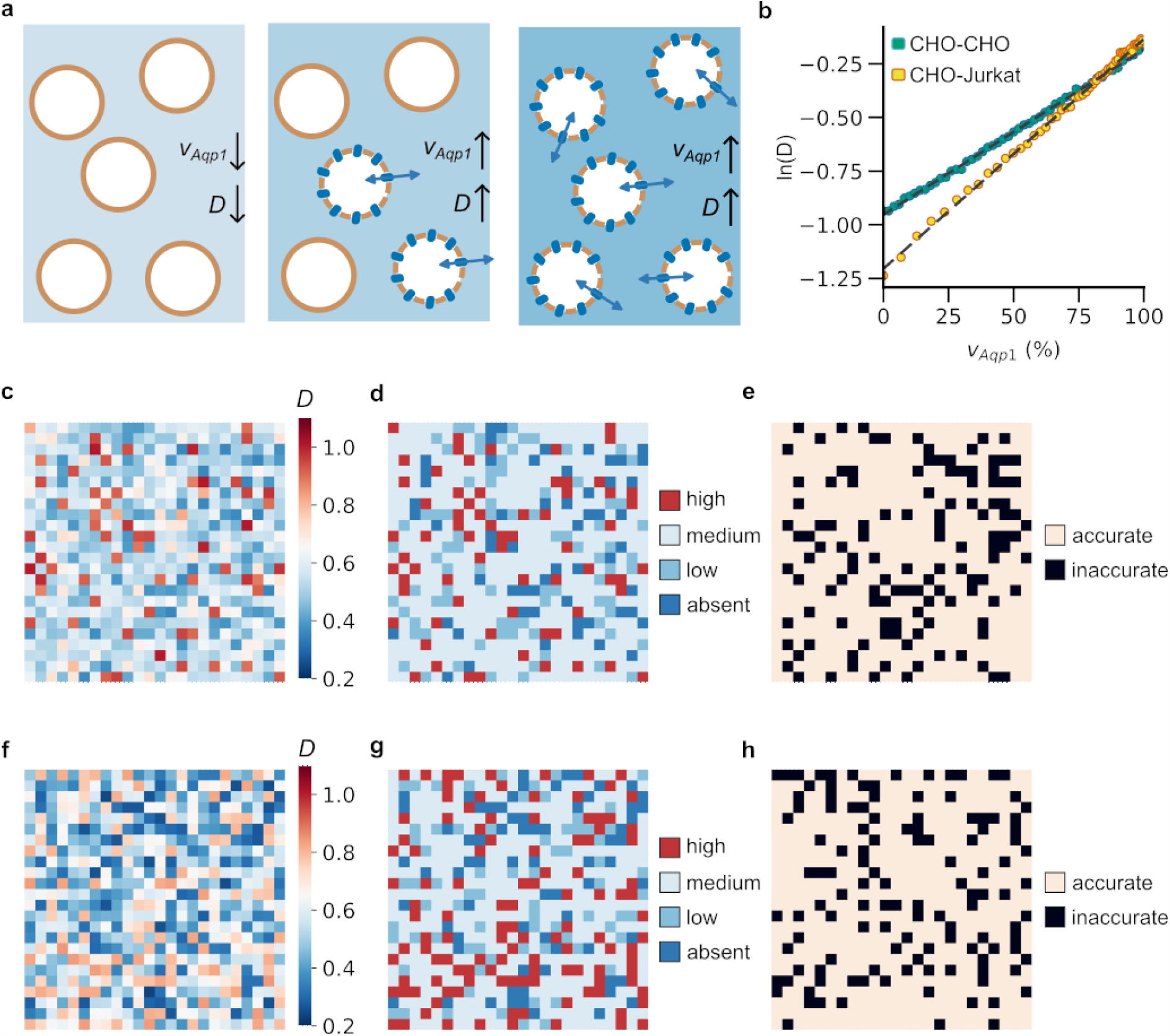
Imaging of mixed-cell populations using Aqp1. **a**, Diffusivity (*D*) of a mixed-cell population increases when more Aqp1-expressing cells are present. **b**, The relationship between diffusivity and volume fraction of Aqp1-expressing cells (*v*_*Aqp*1_) can be modeled by a log-linear function for mixed populations containing Aqp1-expressing CHO cells and either wild-type CHO or wild-type Jurkat cells. **c**, A representative example of a 24 × 24 voxel mixed-cell mosaic created using randomly chosen voxels from experimental diffusion maps of cell populations comprising varying fractions of CHO-Aqp1 cells mixed with wild-type CHO cells. Each voxel in the image corresponds to an experimentally determined diffusion coefficient. **d**, The mapping between diffusivity and *v*_*Aqp*1_ is used to distinguish voxels into one of four levels reflecting the volume percentage of Aqp1-expression: absent (*v*_*Aqp*1_ < 10 %), low (10 % ≤ *v*_*Aqp*1_ < 30 %), medium (30 % ≤ *v*_*Aqp*1_ ≤ 70 %), and high (*v*_*Aqp*1_ > 70 %). **e**, Voxels that are classified correctly are shown in a lighter shade, whereas those that are classified incorrectly are shown in black. **f**, Representative example of a 24 × 24 voxel mixed-cell mosaic created from diffusion maps of mixed populations comprising varying fractions of CHO-Aqp1 cells mixed with wild-type Jurkat cells. **g**, Each voxel was classified into one of four levels, reflecting the volume percentage of Aqp1-expressing cells. **h**, Correctly classified voxels are shown in a lighter shade, whereas those that are classified incorrectly are shown in black. Color-bars in **c, f:** diffusivity (μ*m*^2^/*ms*).

Next, we constructed a mixed-cell mosaic by randomly sampling voxels from experimental diffusion maps of cell populations comprising varying ratios of Aqp1-expressing to wild-type CHO cells (**Fig. 3c**). We used the log-linear mapping between diffusivity and *v*_*Aqp*1_ (**Fig. 3b**) to classify each voxel in the mosaic into one of four groups mirroring the percentage of Aqp1-labeled cells contained in the voxel (**Fig. 3d**). Using this approach, we were able to convert the mixed-cell image into a 4-level classification of Aqp1 volume fraction achieving an accuracy of 79.67 % on entirely unseen experimental data (**Fig. 3e, Table S1**). We further tested this approach on a cell mixture consisting of CHO-Aqp1 cells mixed with a different cell-type, Jurkat T-cells. We simulated Jurkat cells as spheres with a radius of 5 μ*m* and validated that the simulated diffusivity matched with experimental measurements in cell pellets (**Supplementary Fig. 3**). As before, we generated mixed-cell mosaics from diffusion maps of pellets comprising CHO-Aqp1 cells mixed with Jurkat cells in varying ratios (**Fig. 3f**). Finally, we used the log-linear mapping between simulated diffusivity and *v*_*Aqp*1_ for the CHO-Aqp1 and Jurkat mixture (**Fig. 3b**) to perform a 4-level classification of all voxels in the mosaic image, achieving an accuracy of 78.15 % (**Fig. 3g,h, Table S2**).

## DISCUSSION

Here, we quantitatively assessed the performance of Aqp1 as a reporter gene in simulated tissue configurations comprising cells of different radii, volume fractions, and proportions of Aqp1-expressing cells. Our study revealed four major findings that we anticipate will be used to guide the design and analysis of future experiments involving Aqp1 to track cells, genetic function, and molecular activity in living organisms. First, we found that Aqp1 is robust to cell size variations, making it a suitable reporter for cells of different sizes. This prediction is reinforced by a growing body of literature showing that Aqp1 operates as a viable reporter across distinct cell-types, such as tumors, neurons, and glial cells^42–44^. Second, this study emphasizes the importance of accounting for volume fraction when interpreting MRI signals generated by Aqp1, especially when a large reduction in cell density is expected, such as during tumor therapy. Third, we demonstrate that subtracting diffusivities at two time points provides a unique approach for disambiguating Aqp1 signals from the nonspecific effects of tissue microstructure on water diffusion. Finally, we observed that diffusivity was quantitatively linked to the volume fraction of cells expressing Aqp1, suggesting that Aqp1 can be used as a genetic indicator to measure the percentage of reporter-expressing cells in mixed populations.

The current study has limitations, which suggest potential avenues for future research. First, the two-compartment model employed here, similar to those used in past diffusion modeling studies^26,31,34,39^, could be amended to include additional water pools for subcellular structures such as the nucleus and extracellular structures such as the vasculature^27,32,45^. Such multicompartment models could help in analyzing how Aqp1 behaves in systems where nuclear and/or vascular volume fractions vary, which could in turn modify intra- and extracellular diffusion coefficients. Second, our tissue phantoms can be tailored to accurately reflect realistic geometries derived from the histology of engineered tissues expressing Aqp1^46^. Although we expect that the correlations found in this study will hold for even more complex tissue morphologies, histology-derived meshes can be useful for exploring the context-dependent behavior of Aqp1 in realistic in vivo settings. To this end, Monte Carlo diffusion simulations are advantageous due to their ability to integrate new experimental data, which is expected to grow owing to advances in diffusion MRI technology^47^ and the adoption of Aqp1-based reporters by the scientific community^42–44^. Finally, Monte Carlo simulations could be used to train learning algorithms to generate spatial maps of gene expression and cell density based on Aqp1 reporter signals measured in biological tissues^46^. To do so, the diffusion signal must be represented by a more comprehensive feature vector, likely including additional metrics derived from multishell diffusion-weighted imaging experiments^48^. These efforts should be bolstered by machine learning models that use multi-shell diffusion tensor data to compute tissue microstructure parameters^49–52^.

In summary, this study establishes Aqp1 as a quantitative tool for biomedical imaging by bringing together the capabilities of genetic targeting, diffusion biophysics, and Monte Carlo simulations. We envision a rapid expansion of Aqp1 applications in the future, especially with advances in reporter engineering and diffusion imaging technology, synergizing with the computational approach established in this work.

## METHODS

### Monte Carlo diffusion simulations

We modeled cells as packed spheres and dispersed *N* = 10^5^ particles (representing diffusing water spins) evenly in the intra- and extracellular compartments, in proportion to the relative volume of each compartment. We performed two-compartment diffusion simulations with a time step (*τ*) of 19.682 μ*m* using the open-source Monte Carlo Diffusion and Collision Simulator developed in^26^ and later extended to permeable substrates^36^. The total number of particles (*N*) and step size (*τ*) were chosen to ensure accuracy and convergence of the simulation runs^26^. Using these parameters, the standard deviation of simulated diffusivity is less than 2 % of its mean across all simulations. Briefly, at each time step (*τ* = 19 μ*s*), we randomly displaced a spin (*i*) by a distance (*x*_*i*_) computed from Einstein’s diffusion equation assuming diffusion coefficients of 1 µm^2^/ms and 2 µm^2^/ms for the intra- and extracellular compartments, respectively. Upon encountering a cell membrane, a spin can undergo an elastic reflection^25^ or pass through the membrane, with a probability that depends on the permeability coefficient (*κ*) of the membrane and is calculated as described in prior work^46^. At the desired diffusion time (Δ), we sum the total phase dispersion of by all spins in the ensemble to compute the diffusion coefficient (*D*) as follows: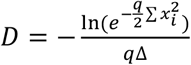.

Here, *q* represents the diffusion-weighting defined as *q* = (*γgδ*)^2^ where *γγ* = 42.57 *MHz*/*T, δ* is the gradient duration, *γγ* is the gradient strength. The values of Δ, *δ*, and *γγ* were based on the experimentally defined diffusion-weighted MRI parameters (see below). We first investigated the change in diffusivity as a function of membrane permeability for a fixed cell radius (*r* = 7.6 μ*m*) and volume fraction (*v*_*f*_ = 0.65). This radius was chosen to be consistent with the effective size of CHO cells^53^ and the volume fraction approximates lightly centrifuged cell pellets as well as many physiological tissues. We investigated three scenarios: (1) *v*_*f*_ = 0.65, 5 μ*m* ≤ *r* ≤ 25 μ*m* (2) *r* = 7.6 μ*m*, 0.10 ≤ *v*_*f*_ ≤ 0.67 (3) concurrently varying both *r* and *v*_*f*_ within the aforementioned limits. For each condition, we tested two permeability coefficients corresponding to wild-type (0.012 μ*m ms*^−1^) and Aqp1-expressing CHO cells (0.138 μ*m ms*^−1^). The permeability values are based on previously published estimates obtained in wild-type CHO cells and CHO cells stably transfected to express Aqp1^40^.

Mixed-cell Monte Carlo simulations were performed in the same manner as described above, but by varying the number of Aqp1-expressing and wild-type CHO cells to achieve a desired Aqp1 volume fraction (*v*_*f,Aqp*1_). For mixed-cell experiments involving CHO and Jurkat cells, the latter were modeled as smaller spheres (*r* = 5 μ*m*) with *κ* = 0.005 μ*m*/*mm*^54,55^. We packed spheres of two different radii (*r*_*CHO*_ = 7.6 μ*m, r*_*Jurkat*_ = 5 μ*m*) to obtain a total intracellular volume fraction of 0.65, as described previously^56^, while adjusting the number of bigger spheres to obtain the desired volume fraction of CHO cells.

### Reagents

Dulbecco’s Modified Eagle Media (DMEM), sodium pyruvate, doxycycline hyclate, and penicillin-streptomycin (10^4^ units/mL penicillin and 10 mg/mL streptomycin) were purchased from Sigma-Aldrich (St. Louis, MO, USA). Roswell Park Memorial Institute media (RPMI 1640), sterile phosphate buffered saline (PBS), TrypLE, and Gibco(tm) fetal bovine serum (FBS) were purchased from Thermo Fisher Scientific (Waltham, MA, USA). MycoAlert® Plus Mycoplasma Detection Assay was purchased from Lonza.

### Cell culture

Chinese hamster ovary (CHO) cell lines were genetically engineered to express human aquaporin-1 (hAqp1) exactly as described in our previous work. Both wild-type and Aqp1-expressing CHO cells were cultured at 37 °C in a humidified 5 % CO_2_ incubator using DMEM supplemented with 10 % FBS, 100 U/mL penicillin, and 100 µg/mL streptomycin. Jurkat cells were grown as suspension culture using RPMI medium supplemented as before.

### In vitro MRI

Approximately 24 h before imaging, cells were treated with doxycycline hyclate (1-10 µg/mL) to activate Aqp1 expression. Adherent CHO cells were harvested by trypsinization, centrifuged at 350 x g, and resuspended in 200 µL sterile PBS in 0.2 mL tubes. For the mixed cell experiments, the two cell types were cultured separately and 10 µL of the cell suspension was loaded in a disposable hemocytometer to count cells using a brightfield microscope. Based on the cell counts, appropriate volumes of the two cell types were mixed to achieve a desired volume fraction of Aqp1-expressing cells. The cells were mixed by gentle pipetting, centrifuged, and transferred to 0.2 mL tubes. The 0.2 mL tubes were centrifuged at a low speed (500 x g for 5 min) to form compact pellets. The pellet-containing tubes were housed in a water-filled agarose (1 % w/v) phantom for imaging. MR images were acquired using a 66 mm diameter coil in a Bruker 7T vertical-bore scanner. Diffusion-weighted images were acquired in the axial plane using a stimulated echo sequence with the following parameters: echo time, T_E_ = 18 ms, repetition time, T_R_ = 1000 ms, gradient duration, δ = 5 ms, gradient separation, Δ = 20, 50, 80, 100, 200, and 300 ms, matrix size = 128 × 128, field of view (FOV) = 5.08 × 5.08 cm^2^, slice thickness = 1-2 mm, number of averages = 5, and 4 nominal b-values: 0, 400, 600, and 800 s/mm^2^. Although the same set of nominal b-values was used at all diffusion times, the effective b-values changed substantially owing to the contribution of imaging gradients to the diffusion weighting via cross terms^57^. Diffusion-weighted intensity was determined using region of interest (ROI) analysis in Fiji (NIH) and the slope of the logarithmic decay in mean intensity versus effective b-value was used to calculate the diffusivity. To generate voxel-wise diffusion maps, a diffusion coefficient was computed for each voxel. Least-squares regression fitting was performed using the fitnlm function in Matlab (R2022b).

### Voxel-wise classification of mixed-cell populations based on Aqp1 volume fraction

We binned individual voxels from experimental diffusion maps (acquired at Δ = 100 ms) of mixed-cell pellets into one of nine groups (0, 10, 20, 30, 40, 50, 60, 80, and 100 %) based on the known Aqp1 volume fraction (*v*_*Aqp*1_) of the pellet. The ensuing dataset comprises approximately 3744 voxels (4 replicates x 104 voxels per image x 9 *v*_*Aqp*1_ values) representing noisy experimental data for a range of Aqp1 volume fractions. We denoised diffusivity values in each bin using a Gaussian filter (*σ* = 3), similar to how experimental diffusion maps are commonly smoothed. From each bin, we sampled 64 voxels with replacement and distributed them randomly in a 24 × 24 grid to construct a “mixed-cell” ADC image. We modeled the dependence of the simulated ADC (Δ = 100 ms) on *v*_*Aqp*1_ using a log-linear function of the form *log*(*ADC*) = *av*_*Aqp*1_ + *b*. We used the resulting log-linear mapping to classify each voxel in the mixed-cell ADC image into one of four levels: absent (*v*_*Aqp*1_ < 10 %), low (10 % ≤ *v*_*Aqp*1_ < 30 %), medium (30 % ≤ *v*_*Aqp*1_ ≤ 70 %), and high (*v*_*Aqp*1_ > 70 %). This discretized 4-level classification was deemed appropriate, given that experimental ADC maps are inherently noisy, particularly at the long diffusion times needed to maximize Aqp1-based contrast. To evaluate the performance of the model, we repeated the 4-level classification on 100 randomly generated sets of mixed-cell images and computed the confusion matrix using the precision_recall_fscore_support package from sklearn.metrics in Python.

### Statistical analysis

Experimental data are summarized by their mean and standard deviation obtained from multiple (*n* ≥ 4) biological replicates defined as measurements performed with distinct cell samples. Quality of model-fitting was judged based on the regression coefficient and inspection of the 95 % confidence intervals and coefficients of determination. All tests are 2-sided and a *P* value of less than 0.05 taken to indicate statistical significance.

## Author Contributions

AM conceived the study. AM and RC designed experiments. RC and JW performed all experiments with insights and feedback from RG, JRP, and JPT. RC and AM performed all data analysis. AM wrote the manuscript with inputs from all authors. AM supervised the research.

## Conflicts of Interest

There are no conflicts to declare.

## Acknowledgements

We thank Dr. Harun F. Ozbakir, Dr. Jason Yun, Dr. Hunter Davis, and Dr. Thomas Oerther (Bruker BioSpin GmbH) for several helpful discussions and optimization of imaging protocols. This research was supported by the National Institutes of Health (R35-GM133530, R03-DA050971, and R01-NS128278 to A.M.), the U.S. Army Research Office via the Institute for Collaborative Biotechnologies cooperative agreement W911NF-19-D-0001-0009 (A.M.), a NARSAD Young Investigator Award from the Brain & Behavior Research Foundation (A.M.), and a UC Santa Barbara Office of Research Early-Stage Seed Grants (A.M. and F.G). This project has also been made possible in part by a grant from the Chan Zuckerberg Initiative DAF, an advised fund of Silicon Valley Community Foundation. All MRI experiments were performed at the Materials Research Laboratory (MRL) at UC, Santa Barbara. The MRL Shared Experimental Facilities are supported by the MRSEC Program of the NSF under Award No. DMR 1720256; a member of the NSF-funded Materials Research Facilities Network. We acknowledge the use of the NRI-MCDB Microscopy Facility and the Resonant Scanning Confocal supported by NSF MRI grant 1625770.

## Notes

### Competing Interest Statement

The authors have declared no competing interest.

## REFERENCES

1. Gilad, A. A. & Shapiro, M. G. Molecular Imaging in Synthetic Biology, and Synthetic Biology in Molecular Imaging. Mol Imaging Biol 19, 373–378 (2017).

2. Schuerle, S. et al. Genetic Encoding of Targeted Magnetic Resonance Imaging Contrast Agents for Tumor Imaging. ACS Synth. Biol. 9, 392–401 (2020).

3. Bricco, A. R. et al. A Genetic Programming Approach to Engineering MRI Reporter Genes. ACS Synth. Biol. 12, 1154–1163 (2023).

4. Helmchen, F. & Denk, W. Deep tissue two-photon microscopy. Nat Methods 2, 932–940 (2005).

5. Mukherjee, A., Davis, H. C., Ramesh, P., Lu, G. J. & Shapiro, M. G. Biomolecular MRI reporters: Evolution of new mechanisms. Progress in Nuclear Magnetic Resonance Spectroscopy 102–103, 32–42 (2017).

6. Mukherjee, A., Wu, D., Davis, H. C. & Shapiro, M. G. Non-invasive imaging using reporter genes altering cellular water permeability. Nat Commun 7, 13891 (2016).

7. Yun, J., Baldini, M., Chowdhury, R. & Mukherjee, A. Designing Protein-Based Probes for Sensing Biological Analytes with Magnetic Resonance Imaging. Analysis &Sensing 2, e202200019 (2022).

8. Benga, G. The first discovered water channel protein, later called aquaporin 1: Molecular characteristics, functions and medical implications. Molecular Aspects of Medicine 33, 518–534 (2012).

9. Verkman, A. S. & Mitra, A. K. Structure and function of aquaporin water channels. American Journal of Physiology-Renal Physiology 278, F13–F28 (2000).

10. Bammer, R. Basic principles of diffusion-weighted imaging. European Journal of Radiology 45, 169–184 (2003).

11. Merboldt, K.-D., Hanicke, W. & Frahm, J. Self-diffusion NMR imaging using stimulated echoes. Journal of Magnetic Resonance (1969) 64, 479–486 (1985).

12. Harkins, K. D., Galons, J.-P., Secomb, T. W. Trouard, T. P. Assessment of the effects of cellular tissue properties on ADC measurements by numerical simulation of water diffusion. Magnetic Resonance in Medicine 62, 1414–1422 (2009).

13. Xu, J., Does, M. D. & Gore, J. C. Quantitative characterization of tissue microstructure with temporal diffusion spectroscopy. Journal of Magnetic Resonance 200, 189–197 (2009).

14. Bihan, D. L. Molecular diffusion, tissue microdynamics and microstructure. NMR in Biomedicine 8, 375–386 (1995).

15. Nilsson, M., van Westen, D., Ståhlberg, F., Sundgren, P. C. &Lätt, J. The role of tissue microstructure and water exchange in biophysical modelling of diffusion in white matter. Magn Reson Mater Phy 26, 345–370 (2013).

16. Pilatus, U. et al. Intracellular volume and apparent diffusion constants of perfused cancer cell cultures, as measured by NMR. Magn Reson Med 37, 825–832 (1997).

17. Norris, D. G. The effects of microscopic tissue parameters on the diffusion weighted magnetic resonance imaging experiment. NMR Biomed 14, 77–93 (2001).

18. Galons, J.-P., Altbach, M. I., Paine-Murrieta, G. D., Taylor, C. W. & Gillies, R. J. Early Increases in Breast Tumor Xenograft Water Mobility in Response to Paclitaxel Therapy Detected by Non-Invasive Diffusion Magnetic Resonance Imaging. Neoplasia 1, 113–117 (1999).

19. Anderson, N. T., Weyant, K. B. & Mukherjee, A. Characterization of flavin binding in oxygen-independent fluorescent reporters. AIChE Journal 66, e17083 (2020).

20. Moffat, B. A. et al. Functional diffusion map: a noninvasive MRI biomarker for early stratification of clinical brain tumor response. Proc Natl Acad Sci U S A 102, 5524–5529 (2005).

21. Jiang, X. et al. Early Detection of Treatment-Induced Mitotic Arrest Using Temporal Diffusion Magnetic Resonance Spectroscopy. Neoplasia 18, 387–397 (2016).

22. Moseley, M. E. et al. Early detection of regional cerebral ischemia in cats: Comparison of diffusion-and T2-weighted MRI and spectroscopy. Magnetic Resonance in Medicine 14, 330–346 (1990).

23. Fieremans, E., Novikov, D. S., Jensen, J. H. & Helpern, J. A. Monte Carlo study of a two-compartment exchange model of diffusion. NMR in Biomedicine 23, 711–724 (2010).

24. Yeh, C.-H. et al. Diffusion Microscopist Simulator: A General Monte Carlo Simulation System for Diffusion Magnetic Resonance Imaging. PLOS ONE 8, e76626 (2013).

25. Szafer, A., Zhong, J. & Gore, J. C. Theoretical Model for Water Diffusion in Tissues. Magnetic Resonance in Medicine 33, 697–712 (1995).

26. Rafael-Patino, J. et al. Robust Monte-Carlo Simulations in Diffusion-MRI: Effect of the Substrate Complexity and Parameter Choice on the Reproducibility of Results. Front Neuroinform 14, 8 (2020).

27. Gilani, N., Malcolm, P. & Johnson, G. An improved model for prostate diffusion incorporating the results of Monte Carlo simulations of diffusion in the cellular compartment. NMR in Biomedicine 30, e3782 (2017).

28. Gilani, N., Malcolm, P. & Johnson, G. A monte carlo study of restricted diffusion: Implications for diffusion MRI of prostate cancer: A Monte Carlo Study of Restricted Diffusion: Implications for Diffusion MRI of Prostate Cancer. Magn. Reson. Med. 77, 1671–1677 (2017).

29. Hall, M. G. & Alexander, D. C. Convergence and parameter choice for Monte-Carlo simulations of diffusion MRI. IEEE Trans Med Imaging 28, 1354–1364 (2009).

30. Karunanithy, G. et al. INDIANA: An in-cell diffusion method to characterize the size, abundance and permeability of cells. J Magn Reson 302, 1–13 (2019).

31. Lee, C.-Y., Bennett, K. M. & Debbins, J. P. Sensitivities of statistical distribution model and diffusion kurtosis model in varying microstructural environments: A Monte Carlo study. Journal of Magnetic Resonance 230, 19–26 (2013).

32. Lee, C.-Y., Bennett, K. M., Debbins, J. P., Choi, I.-Y. & Lee, P. The relationship between diffusion heterogeneity and microstructural changes in high-grade gliomas using Monte Carlo simulations. Magnetic Resonance Imaging 85, 108–120 (2022).

33. Baron, C. A. et al. Reduction of Diffusion-Weighted Imaging Contrast of Acute Ischemic Stroke at Short Diffusion Times. Stroke 46, 2136–2141 (2015).

34. Xing, S. & Levesque, I. R. A simulation study of cell size and volume fraction mapping for tissue with two underlying cell populations using diffusion-weighted MRI. Magn Reson Med 86, 1029–1044 (2021).

35. Tachibana, Y., Duval, T. & Obata, T. Monte Carlo Simulator for Diffusion-weighted Imaging Sequences. Magn Reson Med Sci 20, 222–226 (2021).

36. Gardier, R. et al. Cellular Exchange Imaging (CEXI): Evaluation of a diffusion model including water exchange in cells using numerical phantoms of permeable spheres. Magnetic Resonance in Medicine n/a,.

37. Grussu, F. et al. Diffusion MRI signal cumulants and hepatocyte microstructure at fixed diffusion time: Insights from simulations, 9.4T imaging, and histology. Magnetic Resonance in Medicine 88, 365–379 (2022).

38. Budde, M. D. & Frank, J. A. Neurite beading is sufficient to decrease the apparent diffusion coefficient after ischemic stroke. Proc Natl Acad Sci U S A 107, 14472–14477 (2010).

39. Li, H. et al. Time-Dependent Influence of Cell Membrane Permeability on MR Diffusion Measurements. Magn Reson Med 75, 1927–1934 (2016).

40. Farinas, J., Kneen, M., Moore, M. & Verkman, A. S. Plasma membrane water permeability of cultured cells and epithelia measured by light microscopy with spatial filtering. J Gen Physiol 110, 283–296 (1997).

41. Zhu, F., Tajkhorshid, E. & Schulten, K. Theory and Simulation of Water Permeation in Aquaporin-1. Biophys J 86, 50–57 (2004).

42. Li, M. et al. In vivo imaging of astrocytes in the whole brain with engineered AAVs and diffusion-weighted magnetic resonance imaging. Mol Psychiatry 1–8 (2022) doi:10.1038/s41380-022-01580-0.

43. Zhang, L. et al. Targeting visualization of malignant tumor based on the alteration of DWI signal generated by hTERT promoter–driven AQP1 overexpression. Eur J Nucl Med Mol Imaging 49, 2310–2322 (2022).

44. Zheng, N. et al. A novel technology for in vivo detection of cell type-specific neural connection with AQP1-encoding rAAV2-retro vector and metal-free MRI. NeuroImage 258, 119402 (2022).

45. White, N. S. & Dale, A. M. Distinct effects of nuclear volume fraction and cell diameter on high b-value diffusion MRI contrast in tumors. Magn Reson Med 72, 1435–1443 (2014).

46. Lee, H.-H., Fieremans, E. & Novikov, D. S. Realistic Microstructure Simulator (RMS): Monte Carlo simulations of diffusion in three-dimensional cell segmentations of microscopy images. J Neurosci Methods 350, 109018 (2021).

47. Johnson, G. A. et al. Merged magnetic resonance and light sheet microscopy of the whole mouse brain. Proceedings of the National Academy of Sciences 120, e2218617120 (2023).

48. Caruyer, E., Lenglet, C., Sapiro, G. & Deriche, R. Design of Multishell Sampling Schemes with Uniform Coverage in Diffusion MRI. Magn Reson Med 69, 1534–1540 (2013).

49. Nedjati-Gilani, G. L. et al. Machine learning based compartment models with permeability for white matter microstructure imaging. NeuroImage 150, 119–135 (2017).

50. Palombo, M. et al. New paradigm to assess brain cell morphology by diffusion-weighted MR spectroscopy in vivo. Proceedings of the National Academy of Sciences 113, 6671–6676 (2016).

51. Rensonnet, G. et al. Towards microstructure fingerprinting: Estimation of tissue properties from a dictionary of Monte Carlo diffusion MRI simulations. NeuroImage 184, 964–980 (2019).

52. Nilsson, M. et al. Evaluating the accuracy and precision of a two-compartment Kärger model using Monte Carlo simulations. Journal of Magnetic Resonance 206, 59–67 (2010).

53. Han, Y. et al. Cultivation of Recombinant Chinese hamster ovary cells grown as suspended aggregates in stirred vessels. Journal of Bioscience and Bioengineering 102, 430–435 (2006).

54. Voos, P. et al. Ionizing Radiation Induces Morphological Changes and Immunological Modulation of Jurkat Cells. Frontiers in Immunology 9, p(2018).

55. Ross, P. E., Garber, S. S. & Cahalan, M. D. Membrane chloride conductance and capacitance in Jurkat T lymphocytes during osmotic swelling. Biophys J 66, 169–178 (1994).

56. Skoge, M., Donev, A., Stillinger, F. H. & Torquato, S. Packing hyperspheres in high-dimensional Euclidean spaces. Phys. Rev. E 74, 041127 (2006).

57. Jelescu, I. O. et al. Recommendations and guidelines from the ISMRM Diffusion Study Group for preclinical diffusion MRI: Part 1 --In vivo small-animal imaging. Preprint t https://doi.org/10.48550/arXiv.2209.129 94 (2023).

