## Supplementary Information for "Molecular imaging with aquaporin-based reporter genes: quantitative considerations from Monte Carlo diffusion simulations"

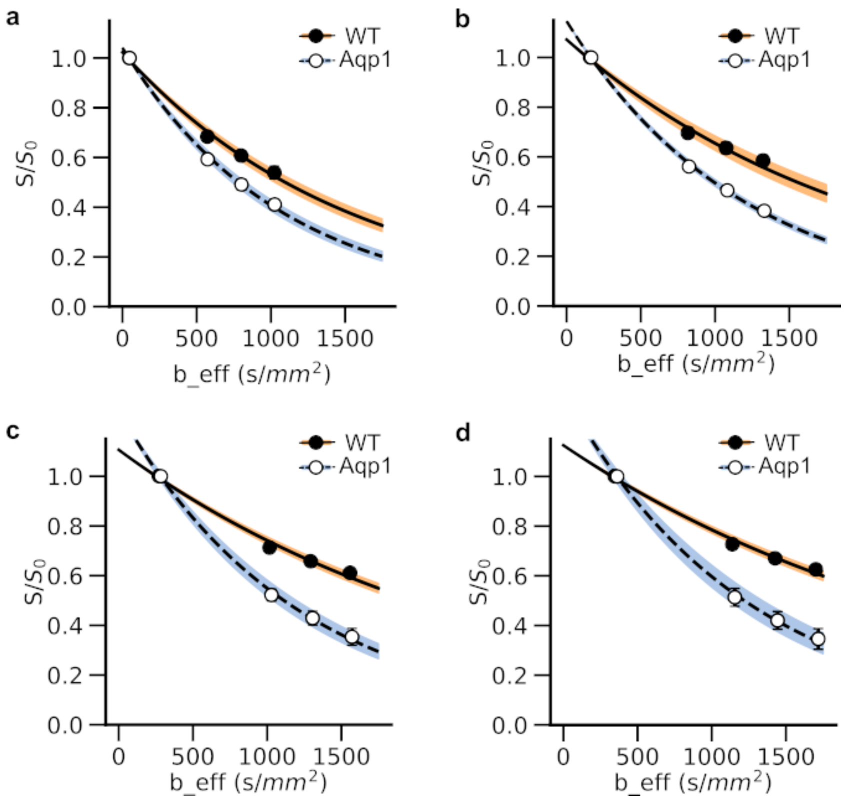


**Figure S1: Measurement of diffusivity in wild-type and Aqp1-expressing cells**. Diffusion-weighted images of pellets of CHO cells were acquired at four effective b-values ($b\_eff$) at various diffusion times (Δ): **a**, 20 ms **b**, 50 ms **c**, 80 ms and **d**, 100 ms. Although the same set of nominal b-values was used at all diffusion times ($b = 0, 400, 600,\ 800\ s/mm^2$), the effective b-values changed substantially owing to the contribution of imaging gradients to the diffusion weighting via cross terms. Accordingly, the effective b-values used at each time are: $46, 572, 800, 1024\ s/mm^2$ (Δ = 20 ms); $158, 817, 1075, 1324\ s/mm^2$ (Δ = 50 ms); $271, 1016, 1293, 1559\ s/mm^2$ (Δ = 80 ms); $345, 1138, 1426, 1701\ s/mm^2$ (Δ = 100 ms). The filled and open circles represent experimental data. The solid and dotted lines represent the mono-exponential fit of the signal decay to the effective b-value. The shaded regions represent 95 % confidence intervals of the fit. Error bars represent standard deviation ($n \geq 5$ biological replicates). All MRI data are acquired at 7 T.

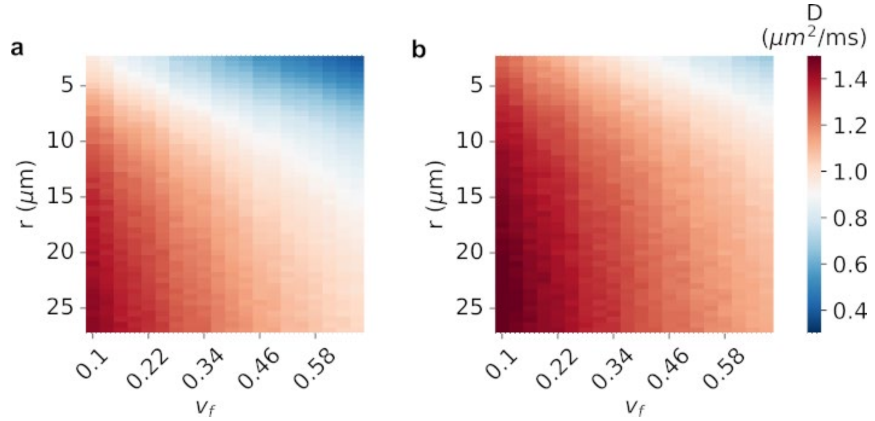


**Figure S2: Effect of cell size and intracellular volume fraction on diffusivity**. Heatmap showing the combined dependence of simulated diffusivity ($D$) on cell radius ($r$) and intracellular volume fraction ($v_f$) in **a**, wild-type and **b**, Aqp1-expressing cells at a diffusion time of 20 ms. Wild-type and Aqp1-expressing cells were modeled with permeability coefficients of $0.012$ and $0.138$ $\mu m/ms$, respectively.

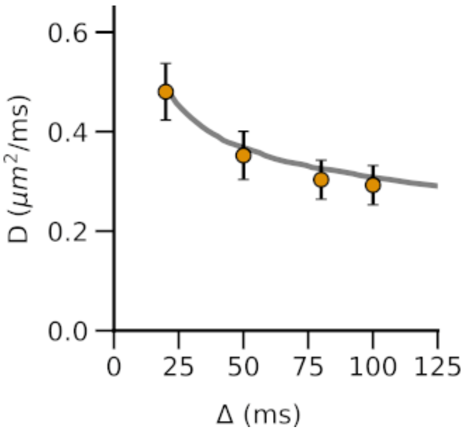


**Figure S3. Monte Carlo simulations of water diffusion in Jurkat cells.** Diffusivity decreases with increasing diffusion time because a larger number of water molecules encounter the plasma membrane, restricting the free movement of water molecules. The solid lines represent the simulated diffusivities for a synthetic substrate consisting of spherical cells of radius $5$ $\mu m$ and permeability $0.005$ $\mu m/ms$ packed to yield a total intracellular volume fraction of $0.65$. Circles denote experimental data obtained from Jurkat cell pellets at 7 T. Error bars represent standard deviation ($n$ = 4 biological replicates).

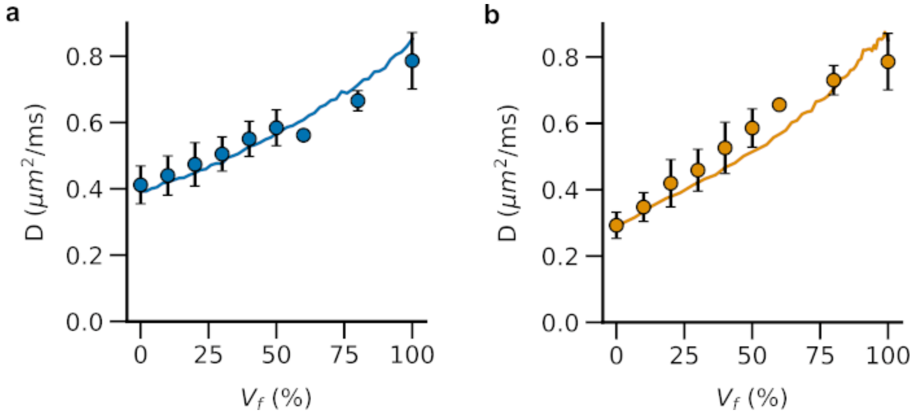


**Figure S4: Imaging of mixed-cell populations using Aqp1**. **a**, Diffusion rate ($D$) of a mixed population increases when more Aqp1-expressing cells are present. **a**, Diffusivities of mixed populations containing Aqp1-expressing CHO cells mixed with wild-type CHO or **b**, wild-type Jurkat cells. Solid line represents the simulated diffusivity. Filled circles denote experimentally measured diffusivity. Error bars represent standard deviation. Wild-type and Aqp1-expressing CHO cells were modeled as spheres of $7.6\,\mu m$ radius with permeability coefficients of $0.012\ \mu m/ms$ and $0.138\ \mu m/ms$ respectively. Wild-type Jurkat cells were modeled as spheres of 5 um radius with a permeability of $0.005\ \mu m/ms$.

**Table S1**: Voxel-wise classification of Aqp1 volume percentage in mixed-cell populations comprising Aqp1-expressing and wild-type CHO cells. Error represents the standard deviation from applying the log-linear classification model on *n* = 100 mixed-cell images constructed by randomly sampling voxels from experimental diffusion maps.

| **Aqp1-level** | **Specificity** | **Recall** | **Precision** |
|---|---|---|---|
| Absent ($<$ 10 %) | 0.953 ± 0.005 | 0.529 ± 0.040 | 0.583 ± 0.031 |
| Low (10 – 30 %) | 0.935 ± 0.006 | 0.383 ± 0.027 | 0.626 ± 0.028 |
| Medium (30 – 70 %) | 0.759 ± 0.014 | 0.975 ± 0.005 | 0.835 ± 0.008 |
| High ($>$ 70 %) | 0.996 ± 0.002 | 1.0 | 0.967 ± 0.017 |

**Table S2**: Voxel-wise classification of Aqp1 volume percentage in mixed-cell populations comprising Aqp1-expressing CHO cells mixed with wild-type Jurkat cells. Error represents the standard deviation from applying the log-linear classification model on *n* = 100 mixed-cell images constructed by randomly sampling voxels from experimental diffusion maps.

| **Aqp1-level** | **Specificity** | **Recall** | **Precision** |
|---|---|---|---|
| Absent ($<$ 10 %) | 0.973 ± 0.003 | 0.885 ± 0.028 | 0.806 ± 0.017 |
| Low (10 – 30 %) | 0.984 ± 0.004 | 0.504 ± 0.020 | 0.898 ± 0.022 |
| Medium (30 – 70 %) | 0.805 ± 0.010 | 0.828 ± 0.006 | 0.842 ± 0.007 |
| High ($>$ 70 %) | 0.892 ± 0.004 | 1.0 | 0.538 ± 0.009 |
